# Astrocytic FMRP regulates the function of spinal parvalbumin-expressing neurons in Fragile X Syndrome

**DOI:** 10.64898/2026.07.17.737291

**Authors:** Haoyi Qiu, Rian Fritz Jalandoni, Ann Derham, Nolan Reinisch, Catherine Chen, Alec Vong, Erik P. Cook, Arjun Krishnaswamy, Anmar Khadra, Reza Sharif-Naeini

## Abstract

Tactile hypersensitivity is a common symptom of Fragile X Syndrome (FXS) characterized by over-responsiveness to innocuous touch or textures. Yet, the neural circuitry underlying this altered sensory processing remains incompletely understood. Previous studies have focused on cortical and peripheral neuron dysfunction. However, the spinal dorsal horn circuits, which make up the first central site of somatosensory integration, also comprises elements that could contribute to the hypersensitivity to innocuous stimuli, yet its involvement in this pathology has remained incompletely understood. Here, we show that spinal parvalbumin (PV)-expressing interneurons (PVNs), an inhibitory population that gates touch sensory input from activating nociceptive pathways, are dysfunctional in FXS. Using a mouse model of FXS (global *Fmr1* knockout; gKO), we showed that dorsal horn PVNs exhibit impaired function characterized by reduced PV expression and inability to sustain tonic firing. To determine whether these deficits arise through cell-intrinsic mechanisms, we selectively deleted *Fmr1* in PVNs. This deletion failed to reproduce the molecular and electrophysiological changes observed in the gKO mice. In contrast, astrocyte-specific deletion of *Fmr1* recapitulated key features of the gKO mice, including decreased PV expression and firing. To understand the biophysical basis of this firing deficit, we simulated spinal PVNs using conductance-based Hodgkin-Huxley type modelling. Surprisingly, modifying intrinsic membrane conductances alone was insufficient to account for the experimental data. In fact, PVN firing required incorporation of an additional calcium-dependent extrinsic synaptic component consistent with the experimental finding that astrocytic, but not PVN-specific, loss of FMRP reproduced the PVN phenotype in gKO mice. Together, these findings show that these deficits in spinal PVNs arise primarily from loss of FMRP in astrocytes rather than in PVNs themselves. It reveals astrocyte-dependent dysfunction of dorsal horn inhibitory circuits as a previously uncharacterized consequence of FXS.

## INTRODUCTION

Fragile X syndrome (FXS) is a neurodevelopmental condition and the leading single gene cause of autism spectrum disorder (ASD). It commonly results from an expansion of a CGG repeat sequence in the *Fmr1* gene that causes hypermethylation, transcriptional silencing, and loss of expression of the fragile X mental retardation protein (FMRP)^1,2^. Through its control of the translation of multiple messenger RNAs^3^, FMRP is critical for synaptic development, circuit maturation, and experience-dependent plasticity^4^. Consequently, loss of FMRP disrupts neural circuit function throughout the nervous system, contributing to the cognitive, social, and sensory abnormalities that characterize FXS^5^.

Among the symptoms experienced by individuals with FXS is tactile hypersensitivity. Patients report aversive responses to otherwise innocuous touch, which can profoundly impair daily activities and social interactions^6,7^. Similar sensory phenotypes are observed in mouse models of FXS, where global *Fmr1* knockout (gKO) mice exhibit enhanced behavioral responses to tactile stimulation^5,8^. Yet the underlying neural circuits responsible for this altered sensory processing remains to be fully elucidated. Studies examining whisker-evoked neuronal activity have demonstrated that cortical circuits are affected by loss of FMRP, with altered excitation-inhibition balance contributing to tactile hypersensitivity^9,10^. Notably, dysfunction of parvalbumin (PV)-expressing interneurons (PVNs) has emerged as a key mechanism underlying cortical sensory abnormalities in FXS, highlighting the contribution of inhibitory circuit maturation for normal sensory perception^11–15^.

Considerably less is known about how tactile information is processed within subcortical sensory pathways in FXS. Previous studies have examined the role of the peripheral nervous system in driving ASD-associated sensory abnormalities, such as tactile aversion and touch hypersensitivity^16,17^, while other work has demonstrated altered cortical excitation-inhibition balance^14,15^. Together, these findings indicate that sensory dysfunction in FXS involves both peripheral and cortical components. However, sensory information is processed sequentially, and the integration of touch input before it reaches the cortex is equally important in determining sensory perception. The spinal dorsal horn represents the first central site where sensory information from peripheral afferents is integrated before being transmitted to ascending pathways and higher-order brain regions. Changes to dorsal horn circuitry could therefore fundamentally reshape touch transmission before they reach higher-order brain regions. Yet whether spinal sensory circuits are disrupted in FXS remains unexplored.

Among dorsal horn inhibitory populations, PVNs are ideally positioned to regulate this transmission by gating low-threshold mechanoreceptor inputs and preventing them from accessing nociceptive pathways. Loss of PVN-mediated inhibition is sufficient to produce mechanical hypersensitivity, implicating these neurons as important regulators of normal touch sensation^18–22^. Given the established involvement of cortical PVNs in FXS^15^ and the presence of hindpaw tactile hypersensitivity in gKO mice^23^, we hypothesized that the function of spinal PVNs may also be disrupted.

Here, we demonstrate that spinal PVNs in gKO mice exhibit impaired function characterized by reduced PV expression and altered intrinsic excitability. Using conditional genetic approaches, we show that these abnormalities are not reproduced by selective deletion of *Fmr1* in PVNs, indicating that cell-intrinsic loss of FMRP is insufficient to account for the phenotype. In contrast, astrocyte-specific deletion of *Fmr1* recapitulates key molecular and electrophysiological features observed in the gKO mice, including reduced PV expression and impaired firing. Finally, computational modeling suggested that alterations in extrinsic synaptic signaling through calcium conductances are required to account for the observed changes in PVN excitability. Together, our findings reveal an astrocyte-dependent mechanism regulating inhibitory circuit function in the spinal dorsal horn.

## RESULTS

### Decreased spinal PVNs in global *Fmr1* KO mice

Alterations in dorsal horn inhibitory circuits have been strongly implicated in pathological sensory states, including chronic pain and tactile hypersensitivity^24–26^. Since gKO mice exhibit tactile hypersensitivity of the hindpaw^23^, we first asked whether inhibitory interneuron populations are altered in the spinal dorsal horn of these animals. Quantification of Pax2-positive inhibitory neurons revealed a reduction in inhibitory neuron count within laminae II–III of gKO mice, whereas no significant change was observed in the superficial dorsal horn (laminae I–II), defined by IB4 labeling^27^ (Fig. S1A–C). These findings suggest that neurons in the deeper dorsal horn laminae are preferentially affected in the gKO mice.

Because PVNs constitute a major inhibitory population within laminae II inner–III and play a critical role in gating tactile sensory input, we examined whether the number of spinal PVNs is altered in the gKO mice. We examined PV immunoreactivity (IR) in the dorsal horn of adult gKO mice. Compared with wildtype (WT), gKO mice have significantly less PV-IR neurons, including inhibitory PVNs identified by Pax2 expression (Fig. 1A–C). Consistent with these findings, *Pvalb* mRNA expression was also lower in the dorsal horn of gKO mice (Fig. S1D–E). However, these observations could reflect either a loss of PVNs or a reduction in PV protein expression within surviving PVNs. To distinguish between these possibilities, we generated a PVcre;tdTomato;*Fmr1* gKO mouse in which PVNs are labeled with tdTomato independent of ongoing PV protein expression. We quantified the temporal expression of tdTomato-positive PVNs across postnatal development at P12, P21, P46, and P60 and found that the number of PVNs and number of inhibitory neurons was significantly lower in PVcre;tdTomato;*Fmr1* gKO mice compared to their WT littermate controls (Fig. 1D–F). Together, these findings indicate that dorsal horn PVNs are consistently lower in developing spinal cord of gKO mice, identifying spinal PVNs as a previously unrecognized cellular population disrupted in FXS.

**Figure 1:**
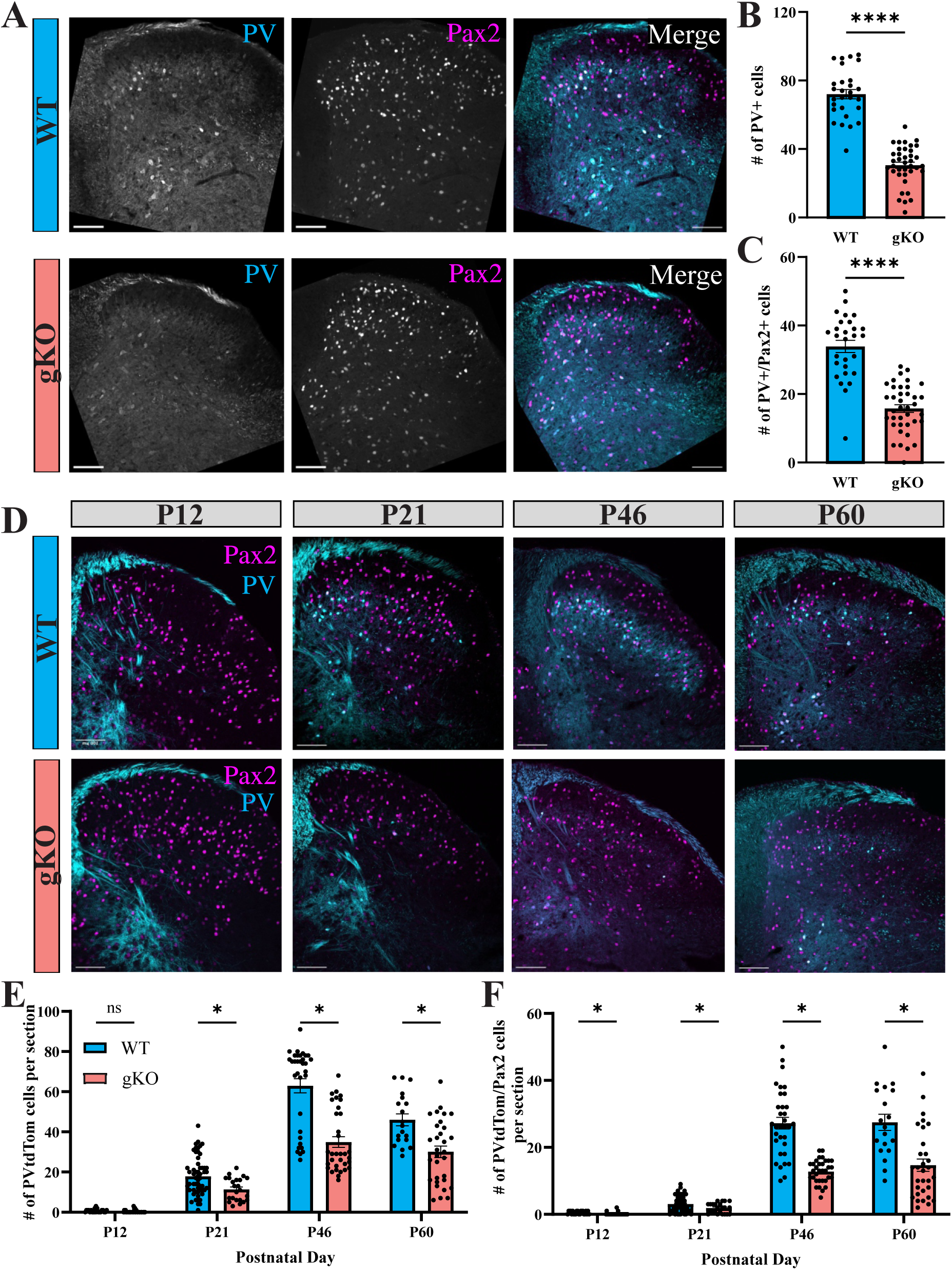
Decreased spinal PVNs in global *Fmr1* KO mice. (A) IHC staining of dorsal horn show total number of cells that showed colocalization between PV-immunoreactivity (IR) and Pax2 to quantify number of inhibitory PVNs in wildtype Fmr1^WT/y^ (WT, top row) and global Fmr1^KO/y^ mice (gKO, bottom row); scale bar = 100 μm. (B-C) Mean ± S.E.M of the number of PV-IR+ (B) and PV-IR+/Pax2+ (C) colocalized neurons in A. WT: n = 28 sections from 4 male mice; gKO: n = 38 sections from 5 male mice, unpaired t-test, two-tailed. ****p-value < 0.0001. (D) Endogenous PV fluorescence (cyan) and IHC staining of Pax2 (magenta) in wildtype littermate PVcre;tdTomato; Fmr1^WT/y^ (WT, top row) and PVcre;tdTomato; global Fmr1^KO/y^ (gKO, bottom row) mice aged postnatal day 12, 21, 46, or 60; scale bar = 100 μm. (E-F) Mean ± S.E.M of the number of PV-tdtomato+ (E) and PV-tdtomato+/Pax2+ (F) colocalized neurons in D. P12: WT = 22 sections from 2 mice, gKO = 34 sections from 3 mice; P21: WT =58 sections from 5 mice, gKO = 23 sections from 2 mice; P46: WT = 33 sections from 3 mice, gKO = 33 sections from 3 mice; P60: WT = 19 sections from 2 mice, gKO =31 sections from 3 mice. Unpaired t-test, two-tailed. *p-value < 0.05.

### Spinal PVNs have decreased intrinsic excitability in global *Fmr1* KO mice

We examined whether these molecular changes observed in PVNs of gKO mice were accompanied by functional deficits. PVNs in multiple regions of the central nervous system are classically characterized as fast-spiking interneurons capable of sustaining high-frequency firing with minimal adaptation^22,28^. Therefore, we examined whether loss of FMRP alters the intrinsic firing properties of spinal PVNs. Using targeted whole-cell patch-clamp recordings from tdTomato-labeled PVNs in PVcre;tdTomato;*Fmr1* gKO mice and WT littermate controls, we assessed intrinsic firing during prolonged depolarizing current injections. In response to a 1-s, 200-pA current step, PVNs from WT mice displayed non-adaptive firing, whereas PVNs from gKO mice exhibited spike-frequency adaptation and a progressive reduction in firing throughout the current injection period (Figure 2A–D). Despite these differences in firing behavior, action potential properties, including spike threshold, amplitude, half-width, and phase-plane characteristics, were similar between genotypes (Figure 2A–B; Supplementary Table 1), suggesting that the fundamental mechanisms underlying action potential generation remained intact. Further analyses of the spike train revealed that PVNs from gKO mice exhibited a significantly reduction in frequency decay time constant (τ_f_), a shorter duration of sustained spiking (t_d_), an increased adaptation index, and a lower total spike count during the current injection (Figure 2D-E). These findings indicate that loss of *Fmr1* does not substantially alter action potential generation itself but instead reduces the ability of PVNs to maintain repetitive firing, resulting in spike-frequency adaptation.

**Figure 2:**
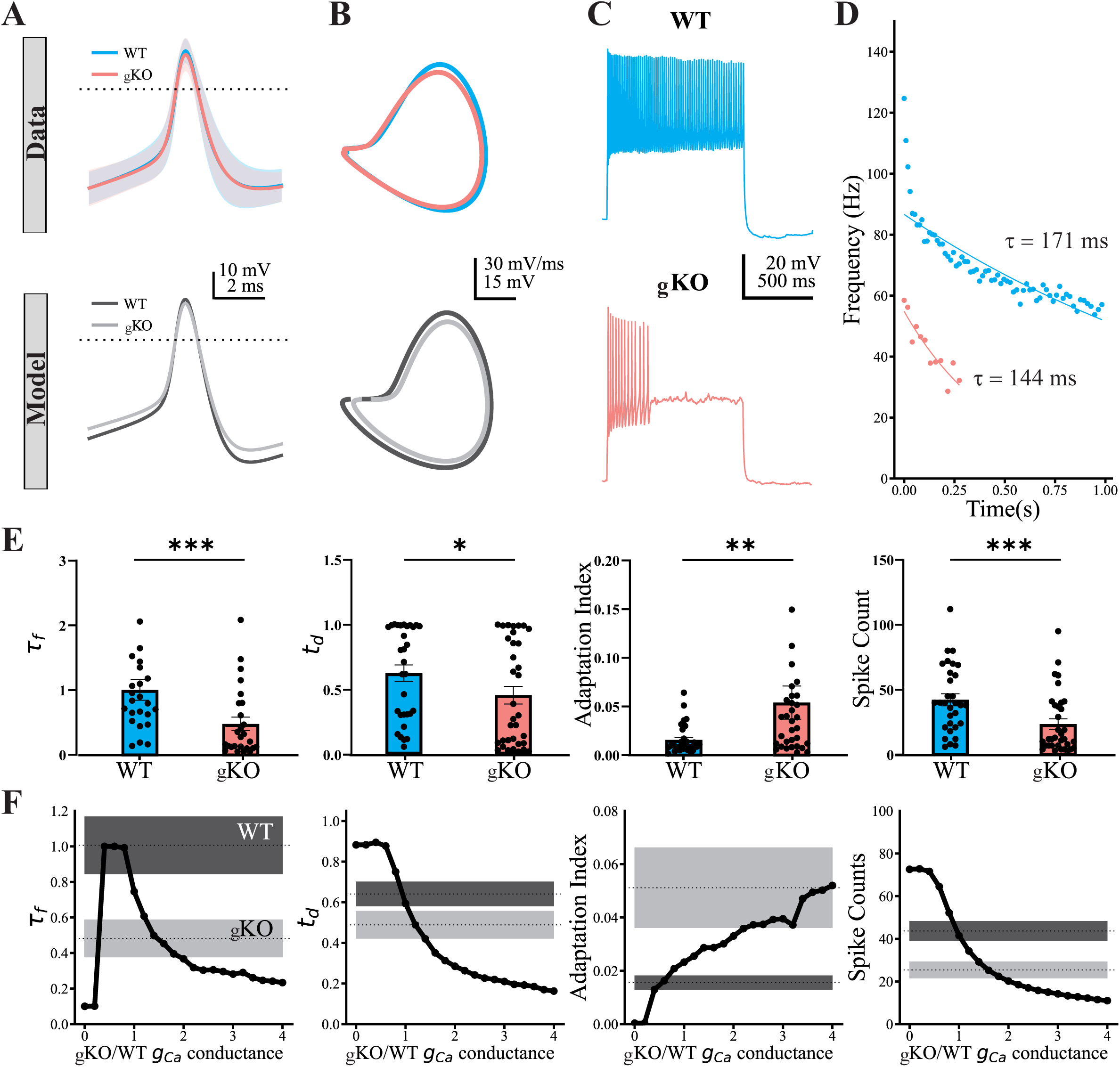
Spinal PVNs have decreased intrinsic excitability in global *Fmr1* KO mice. (A) Mean (dark) ± S.E.M (shaded) traces of action potentials of dorsal horn PVNs (top row) and its simulated recordings from the computational model (bottom row) in wildtype littermate PVcre;tdTomato; Fmr1^WT/y^ (WT, black) and PVcre;tdTomato; global Fmr1^KO/y^ (gKO, grey). (B) Phase plot of the action potentials in A (top row) and the simulation from the computational model (bottom row). (C) Representative spiking trace of a PVN in response to a 1-second 200 pA step current injection in WT and gKO mice. (D) Instantaneous spike frequencies (dots) of a PVN from WT (black) and gKO (grey) in C measured as the inverse of interspike intervals and fitted to a single exponential decay function (lines). (E) Mean ± S.E.M of the frequency decay constant (τ_f_), discharge time (*t_d_*), Adaptation index and spike count in PVNs from WT (blue bar) and gKO (red bar) mice. n= 31 cells from 5 WT mice, n= 34 cells from 8 gKO mice. (F) Average changes in frequency decay constant, discharge time, adaptation index, and spike count across a heterogeneous population of model cells (n=30) as calcium (Ca^2+^) conductance density was progressively increased in the simulated gKO condition, relative to the WT baseline conductance density. The experimental mean (shaded) ± S.E.M (dotted line) WT and gKO ranges are indicated in dark and light grey respectively.

To gain mechanistic insights into the ionic processes that could underlie this firing phenotype, we next developed a conductance-based Hodgkin–Huxley (HH) model of spinal PVNs adapted from previous published experimentally constrained PVN models ^20,29,30^. Model parameters were re-optimized to reproduce the firing behavior and phase-plane trajectories observed in WT PVNs (Figure 2A), generating a baseline WT-PVN model that accurately captured both action potential waveform dynamics, spike train features, and the voltage phase-plane plots (Table 1) using the population-based parameter fitting method (see Methods). Fitting the phase-plane trajectories (V, dV/dt) in addition to the spike waveform was essential because the waveform alone can be reproduced by many different combinations of ionic conductances, leaving a bigger space of parameter degeneracy ^31^. Because dV/dt is directly proportional to the net membrane current, the phase-plane trajectory constrains the kinetics and relative magnitudes of the underlying sodium and potassium currents. This addition ensures that the model reproduces not only the observed firing but also the ionic dynamics that generate it, which is a prerequisite for reliably predicting how the phenotype changes under perturbation. If a single conductance underlies the firing differences between WT and gKO PVNs, then systematically varying that conductance *in silico* should be sufficient to shift the measured firing parameters in Figure 2E from the WT regime into the gKO regime. Increasing voltage-gated calcium conductance at the population level progressively shifted model firing patterns toward the gKO phenotype, reducing τ_f_, t_d_, and total spike count, while increasing the adaptation index (Figure 2F), encompassing the experimentally observed differences. Perturbations of other conductances failed to recapitulate the full phenotype changes in gKO (Figure S2). Together, these results demonstrate that spinal PVNs in gKO mice exhibit impaired firing characterized by frequency adaptation, and computational modeling identifies altered calcium-dependent signaling as the most plausible mechanism underlying this change.

**Table 1.**
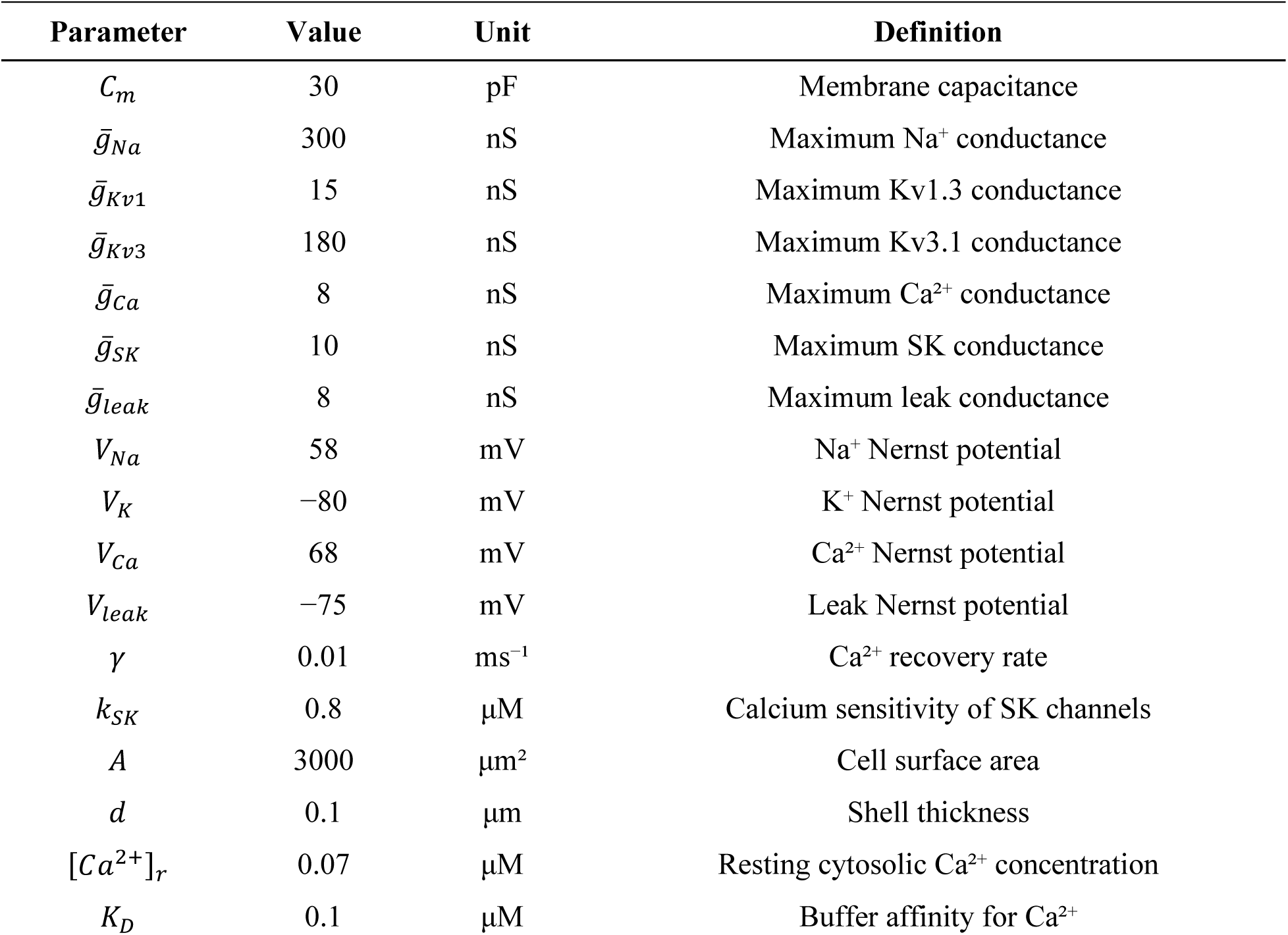
PVIN model parameters.

### *Fmr1* deletion in PVNs does not account for deficits in global *Fmr1* KO mice

The widespread expression of FMRP throughout the nervous system raises the question of whether the cellular and functional deficits of spinal PVNs are caused *intrinsically,* by the loss of *Fmr1* within PVNs themselves, or *extrinsically*, through deficits of *Fmr1* deletion in other cell types. Thus, we examined whether selective deletion of *Fmr1* in PVNs was sufficient to reproduce the molecular and functional deficits observed in the spinal cord of gKO mice. To do so, we generated PVcre;tdTomato *Fmr1* conditional knockout (cKO) mice (Figure 3A). Single-cell qPCR performed on tdTomato-labeled PVNs confirmed deletion of *Fmr1*, with significantly lower *Fmr1* expression in PVNs from cKO mice compared with WT littermate controls (Figure 3B). Despite this reduction, *Pvalb* mRNA expression was unchanged between genotypes (Figure 3C). We next examined whether the number of PVNs was affected by cell-intrinsic deletion of *Fmr1*. Quantification of tdTomato-positive PVNs revealed no difference in the total number of PVNs or inhibitory PVNs between cKO and WT littermate mice (Figure 3D–F). These findings contrast sharply with the reduced number of PVNs observed in gKO mice and indicate that selective loss of *Fmr1* in PVNs is insufficient to reproduce the cellular deficits associated with global FMRP loss in the spinal cord.

**Figure 3:**
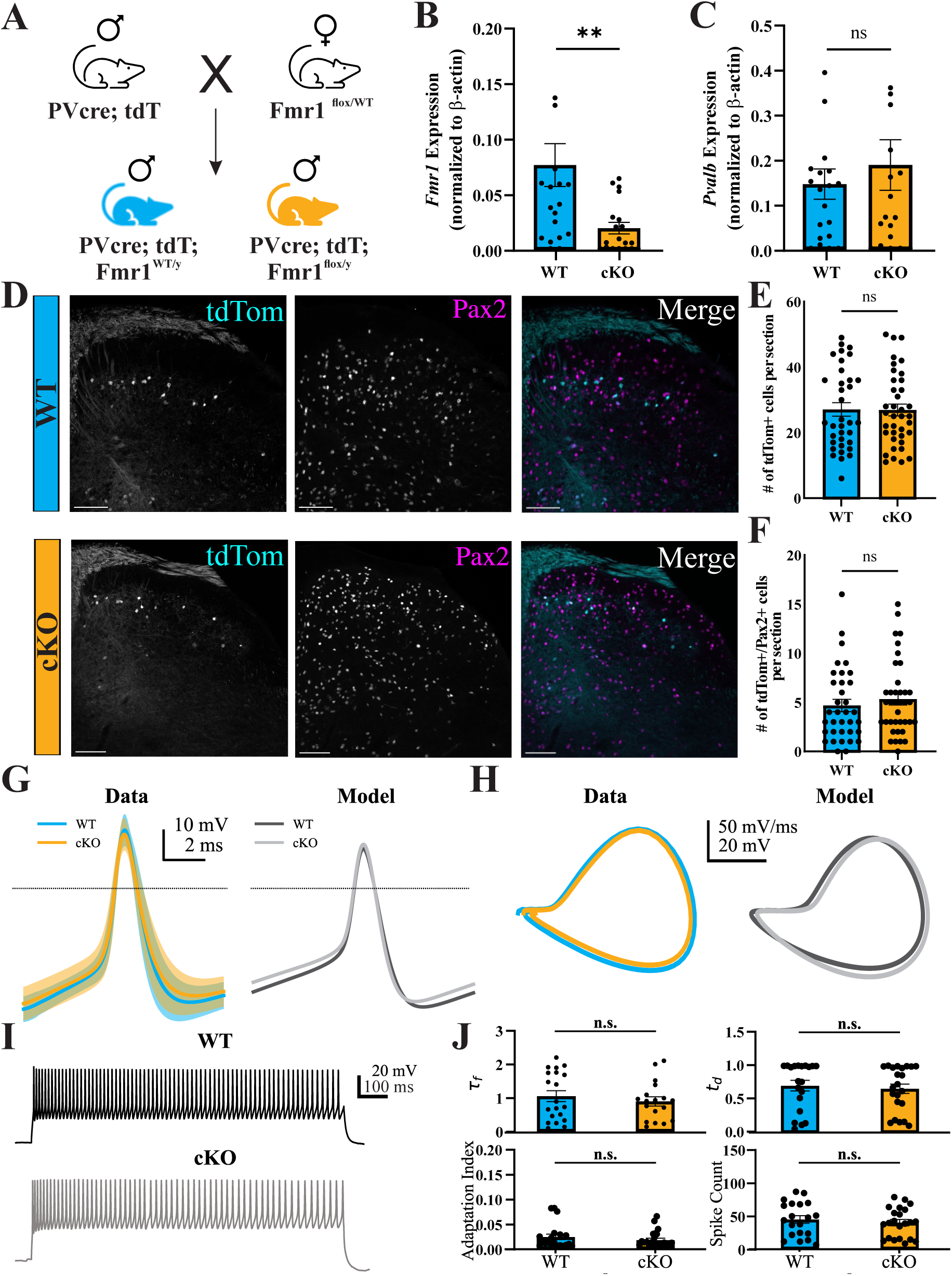
*Fmr1* deletion in PVNs does not account for deficits in spinal PVNs of global *Fmr1* KO mice. (A) Schematic diagram of the breeding scheme to generate male PVcre;tdtomato;Fmr1^flox/y^ mice (cKO, yellow) and littermate wildtype controls (WT, blue). (B-C) Mean ± S.E.M of *Fmr1* (B) and *Pvalb* (C) from single cell-qPCR of PVNs in WT and cKO mice. n= 20 cells from 3 WT mice, n= 19 cells from 4 cKO mice. Unpaired t-test, two-tailed, **p-value < 0.01. (D) Endogenous PV fluorescence (cyan) and IHC staining of Pax2 (magenta) in PVcre;tdTomato;Fmr1^WT/y^ (top row) and PVcre;tdTomato;Fmr1^flox/y^ (bottom row) mice; scale bar = 100 μm. (E-F) Mean ± S.E.M of the number of PV-tdtomato+ (E) and PV-tdtomato+/Pax2+ (F) colocalized neurons in D. n= 35 sections from 4 WT mice, n= 38 sections from 4 cKO mice. Unpaired t-test, two-tailed. (G) Mean (dark) ± S.E.M (shaded) traces of action potentials of dorsal horn PVNs (left) and its simulated recordings from the computational model (right) in PVcre;tdTomato;Fmr1^WT/y^ (blue) and PVcre;tdTomato;Fmr1^flox/y^ (yellow). (H) Phase plot of the action potentials in G (left) and the simulation from the computational model (right). (I) Representative spiking trace of a PVN in response to a 1-second 200 pA step current injection in WT and cKO mice. (J) Mean ± S.E.M of the frequency decay constant, discharge time, adaptation index and spike count in PVNs from WT (blue bar) and cKO (yellow bar) mice. n= 20 cells from 3 WT mice, n= 21 cells from 4 cKO mice.

Next, we asked whether PVN-specific deletion of *Fmr1* altered neuronal function. Targeted whole-cell recordings from tdTomato-labeled PVNs revealed no significant differences in action potential properties between cKO and WT mice (Figure 3G–H; Supplementary Table 2). We generated computational models for PVNs of cKO and WT littermate mice using the same population-based fitting pipeline as in the gKO case. Consistent with the recordings, the cKO and WT simulations showed no significant differences in action potential waveform or phase-plane trajectories (Figure 3G-H). Furthermore, spike train properties that were altered in gKO mice were unchanged between cKO and WT genotypes (Figure 3I–J). Together, these results demonstrate that cell-intrinsic loss of *Fmr1* in PVNs is not sufficient to reproduce either the cellular or functional differences observed in spinal PVNs of gKO mice. These findings suggest that the molecular and functional defects of PVNs arise through non-cell-intrinsic mechanisms and implicate extrinsic components of the dorsal horn circuitry.

### *Fmr1* deletion in astrocytes partly accounts for deficits observed in global *Fmr1* KO mice

Glial cells have emerged as important regulators of neuronal maturation, synaptic function, and inhibitory circuit development^32–35^, and recent studies have implicated microglial, astrocytic, and oligodendrocytic loss of FMRP in the pathophysiology of FXS^36,37^. Therefore, we asked whether dysfunction of glial populations could account for the molecular and functional deficits observed in spinal PVNs. Among glial cell types, astrocytes represent particularly strong candidates because of their established roles in regulating inhibitory neuron maturation and synaptic transmission^33^. Consistent with potential astrocyte involvement, we observed a significantly lower GFAP immunofluorescence intensity in lamina I-III of the dorsal horn of gKO mice despite no change in total GFAP-positive area (Figure S3A–C), suggesting altered astrocyte molecular properties without changes in astrocyte territorial coverage.

Recent work further demonstrated that astrocyte-specific deletion of *Fmr1* reduces PV expression in cortical interneurons, raising the possibility that astrocytic dysfunction may similarly contribute to the deficits observed in spinal PVNs. Therefore, we tested whether specific deletion of *Fmr1* in astrocytes is sufficient to reproduce the molecular and functional deficits observed in gKO mice. To this end, we generated Aldh1l1CreERT2;Pvalb-tdTomato;Fmr1 cKO mice (hereafter referred to as astrocyte-KO, aKO), in which tamoxifen administration induces Cre recombination selectively in astrocytes while tdTomato labels PVNs independently of Cre expression (Figure 4A). Tamoxifen was administered at postnatal days P10–P12, prior to the emergence of tdTomato-positive PVNs within laminae II–III of the dorsal horn (Figure 1D), allowing us to assess the contribution of astrocytic FMRP during PVN development. Immunohistochemical analyses confirmed a significant reduction of FMRP expression within GFAP-positive astrocytes in aKO mice, while total GFAP-positive area remained unchanged (Figure 4B–D), validating astrocyte-specific deletion. Next, we examined whether loss of astrocytic FMRP impacts the number of PVNs. Quantification of tdTomato-positive neurons revealed no difference in the total number of PVNs or inhibitory PVNs between aKO mice and WT littermate controls (Figure 4E–G). However, analysis of tdTomato fluorescence intensity, which reflects activity of the endogenous *Pvalb* promoter in this reporter line, revealed a significant reduction in PV levels within individual PVNs of the aKO mice (Figure 4H–I). Thus, astrocyte-specific deletion of *Fmr1* reduces PV expression without altering PVN survival.

**Figure 4:**
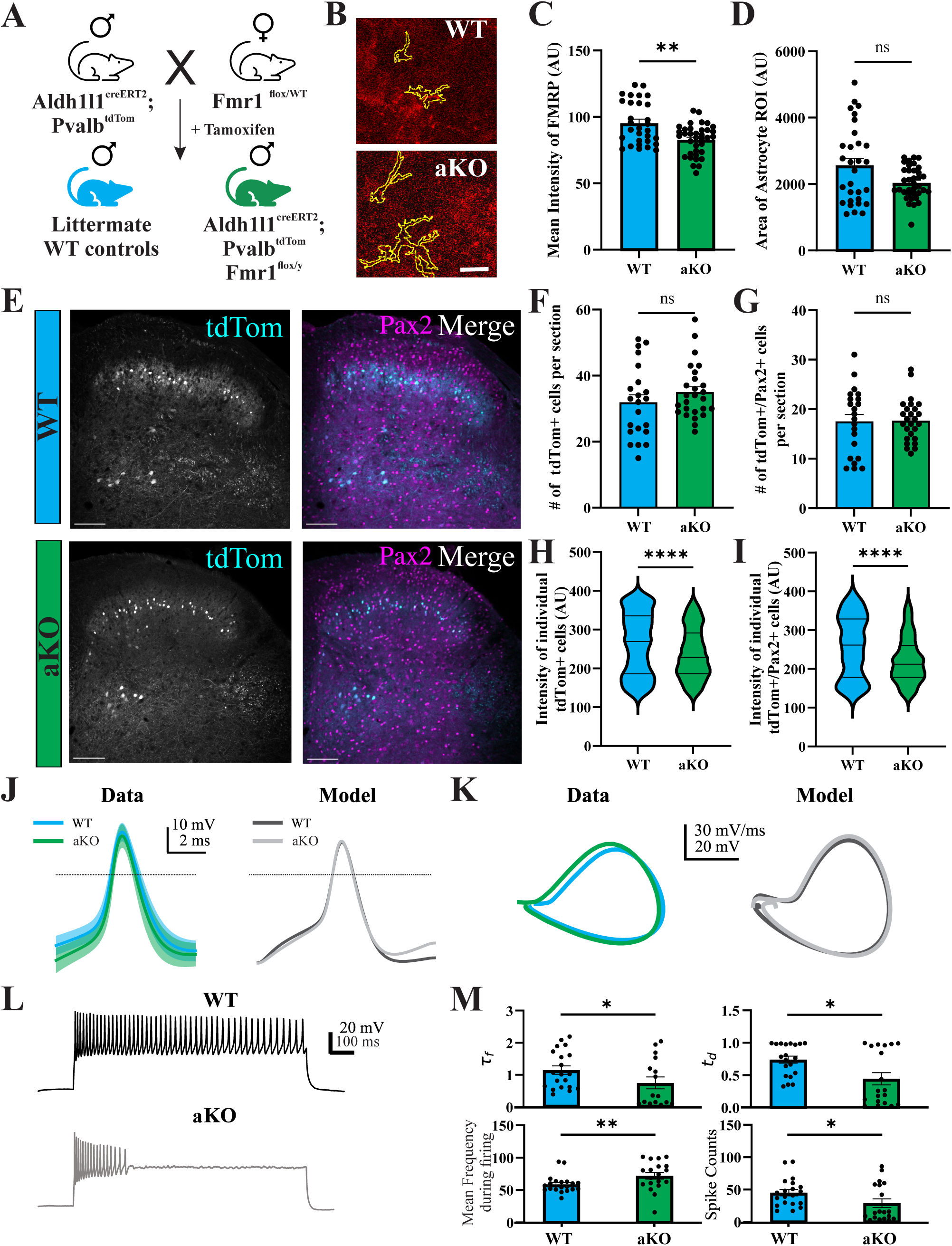
*Fmr1* deletion in astrocytes partly accounts for deficits in spinal PVNs of global *Fmr1* KO mice. (A) Schematic diagram of the breeding scheme to generate male Aldh1l1^CreERT2^;Pvalb^tdTomato^; Fmr1^cKO/y^ mice (aKO, green) and their littermate WT controls (WT, blue). (B-D) Representative images (B), mean ± S.E.M of the mean intensity of FMRP in GFAP-positive astrocytes (C), and the area of the GFAP-positive astrocytes (D) in WT and aKO mice. Yellow outline shows the GFAP-positive region of interests selected for FMRP intensity quantification. n= 30 dorsal horns from 3 WT mice, n= 36 dorsal horns from 3 aKO mice. Unpaired t-test, two-tailed, **p-value = 0.0061. scale bar = 5 μm. (E) Endogenous PV tdTomato fluorescence (cyan) and IHC staining of Pax2 (magenta) in WT (top row) and aKO (bottom row) mice; scale bar = 100 μm. (F-G) Mean ± S.E.M of the number of PV-tdtomato+ (F) and PV-tdtomato+/Pax2+ (G) colocalized neurons in E. n= 22 sections from 3 WT mice, n= 26 sections from 3 aKO mice. Unpaired t-test, two-tailed. (H-I) Mean ± S.E.M of the intensity of tdTomato fluorescence in individual PV-tdtomato+ (H) and PV-tdtomato+/Pax2+ (I) colocalized neurons in E. n= 22 sections from 3 WT mice, n= 26 sections from 3 aKO mice. Unpaired t-test, two-tailed. ****p-value < 0.0001. (J) Mean (dark) ± S.E.M (shaded) traces of action potentials of dorsal horn PVNs (left) and its simulated recordings from the computational model (right) in WT (blue) and aKO (green). (K) Phase plot of the action potentials in J (left) and the simulation from the computational model (right). (L) Representative spiking trace of a PVN in response to a 1-second 200 pA step current injection in WT and aKO mice. (M) Mean ± S.E.M of the frequency decay constant, discharge time, mean frequency, and spike count in PVNs from WT (blue bar) and aKO (green bar) mice. n= 20 cells from 4 WT mice, n= 19 cells from 4 aKO mice.

Previous studies have shown that PV expression is correlated with PVN activity^38–40^. Thus, we asked whether these molecular changes were accompanied by changed in PVN excitability. Targeted whole-cell recordings from tdTomato-labeled PVNs revealed no significant differences in action potential properties between aKO and WT littermate controls (Figure 4J–K; Supplementary Table 3). We fitted separate models for aKO and WT PVNs using the same population-based fitting pipeline as before, now incorporating AMPA and NMDA synaptic currents to capture astrocyte-dependent modulation of PVN input (see Methods). Consistent with the recordings, the aKO and WT models showed no significant differences in action potential waveform or phase-plane trajectories (Figure 4J-K). In contrast, analyses of spiking properties revealed significant differences. PVNs from aKO mice exhibited spike-frequency adaptation, reflected by a reduction in τ_f_, shorter t_d_, and decreased total spike count during sustained 1-second 200 pA step current injection (Figure 4L–M). Notably, these changes closely resembled the firing deficits observed in PVNs from gKO mice (Figure 2E). However, not all parameters of the gKO phenotype were recapitulated in the aKO genotype. Unlike PVNs from gKO mice, PVNs from aKO mice did not exhibit an increase in adaptation index and instead displayed an increase in mean frequency during firing (Figure 4M). Together, these findings demonstrate that astrocyte-specific deletion of *Fmr1* is sufficient to reproduce key molecular and functional features of the PVN phenotype observed in gKO mice. While astrocytic loss of FMRP does not fully account for all aspects of PVN dysfunction, it is sufficient to recapitulate reductions in PV expression and impair the ability of PVNs to sustain high frequency firing, identifying astrocytes as an important non-cell-intrinsic contributor to PVN function in FXS.

### *Fmr1* deletion in astrocytes prolongs spontaneous excitatory synaptic events onto PVNs

Our previous computational analyses focused exclusively on intrinsic membrane conductances and did not account for extrinsic synaptic inputs. Given the established role of astrocytes in regulating synaptic transmission^32,41^ and the partial recapitulation of the gKO phenotype in aKO mice, we next asked whether astrocytic loss of FMRP alters excitatory synaptic input onto PVNs. To address this question, we recorded spontaneous excitatory postsynaptic currents (sEPSCs) from spinal PVNs in aKO mice and WT littermate controls. For each recording, sEPSCs were detrended and low-pass filtered using a zero-phase 4^th^-order Butterworth filter with a cutoff frequency of 2.5 kHz to facilitate event detection and waveform analysis. Peak-aligned averaging of individual sEPSC events revealed clear differences between genotypes (Figure 5A). Neither event frequency nor holding current differed between genotypes, indicating that overall excitatory input rate and baseline membrane conditions were unchanged (Figure 5B-C). In contrast, sEPSC peak amplitude, decay time, and total charge transfer were all significantly increased in aKO PVNs (Figure 5D-F), suggesting stronger and more prolonged excitatory synaptic signaling. These findings demonstrated that astrocyte-specific loss of FMRP selectively enhances the strength and duration of excitatory synaptic events without altering their frequency, resulting in prolonged excitatory drive onto spinal PVNs.

**Figure 5:**
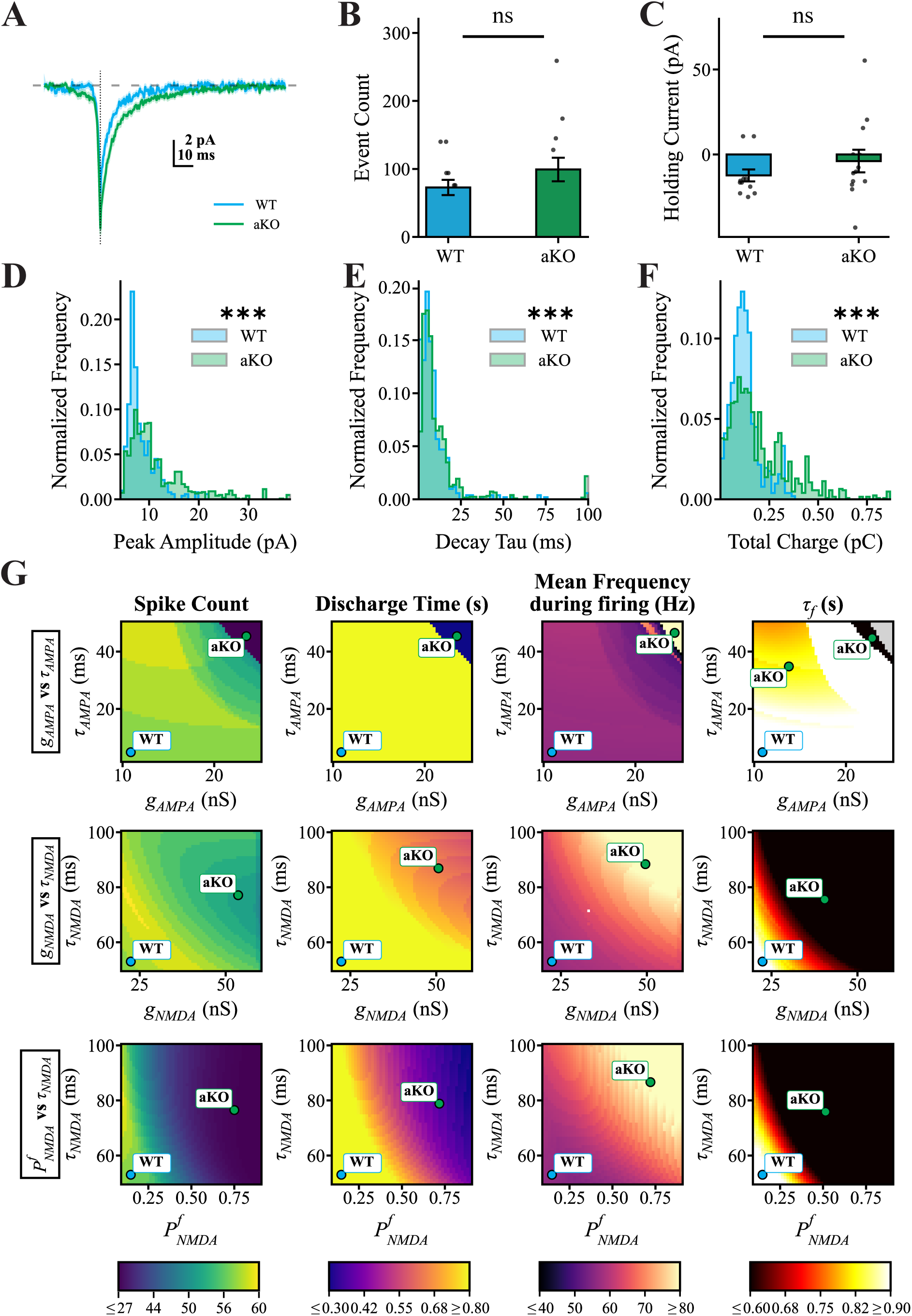
Fmr1 deletion in astrocytes prolongs spontaneous synaptic events in PVNs. (A) Average spontaneous excitatory post-synaptic currents (sEPSCs) in PVNs from Aldh1l1^CreERT2^;Pvalb^tdTomato^; Fmr1^cKO/y^ mice (aKO, green) and their littermate WT controls (WT, blue). (B–F) The event count, holding current, peak amplitude distribution, decay tau distribution, and total charge distribution of the sEPSCs in PVNs from WT and aKO mice. n = 11 cells from 3 WT mice, n = 13 cells from 4 aKO mice. Event count per cell, WT: 72.75 ± 11.25 vs aKO: 99.15 ± 17.38, Mann-Whitney U test, p-value = 0.264, ns. Holding current, WT: −12.30 ± 3.52 pA vs aKO: −3.88 ± 6.64 pA, Mann-Whitney U test, p-value = 0.201, ns. Peak amplitude distribution (WT: n = 511 events; aKO: n = 1236 events), Kolmogorov-Smirnov test, D = 0.294, ***p-value < 0.001. Decay tau distribution (WT: n = 511 events; aKO: n = 1236 events), Kolmogorov-Smirnov test, D = 0.113, ***p-value < 0.001. Total charge distribution (WT: n = 511 events; aKO: n = 1236 events), Kolmogorov-Smirnov test, D = 0.265, ***p-value < 0.001. Values are shown as mean ± S.E.M. (G) Parameter sweep analyses for synaptic current properties in the form of heatmaps showing a progressive shift from WT to aKO-like spike train properties based on AMPA decay time vs. conductance (top row), NMDA decay time vs conductance (middle row), and NMDA decay time vs. calcium permeability defined by the fractions of the NMDA-mediated current carried by calcium 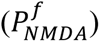 (bottom row). Heatmaps are color-coded according to color-bars at the bottom. Blue dots denote the WT baseline parameter values, and green dots indicate the parameter combinations that is most consistent with the mean spike-train features measured in aKO phenotype.

We next asked whether these synaptic alterations could account for the firing abnormalities observed in aKO PVNs. To test this possibility, we extended our conductance-based PVN model to incorporate both AMPA- and NMDA-mediated synaptic currents (Figure 4J-K; see Methods and Table 2). If prolonged excitatory synaptic signaling contributes to the aKO firing phenotype, then increasing AMPA- and/or NMDA-mediated conductance and decay kinetics in the model should reproduce the pattern of firing changes observed experimentally in aKO PVNs. Baseline synaptic parameters were optimized to reproduce the firing properties of WT PVNs, after which synaptic conductance and decay kinetics were systematically varied to mimic the changes observed experimentally. Increasing AMPA conductance and decay time constant produced a reduction in spike count and discharge duration, accompanied by a decrease in frequency decay time constant and an increase in mean firing frequency (Figure 5G, top panel). Notably, these changes matched the firing phenotype observed in aKO PVNs. Similar effects emerged with increases in NMDA conductance and decay kinetics (Figure 5G, middle panel), but the influence was weaker, yielding higher spike counts and longer discharge times than those produced by AMPA perturbations. Interestingly, our previous simulations showed that increasing intrinsic calcium conductance was able to reproduce several features of the gKO PVNs (Figure 2G). Because astrocyte-specific deletion of *Fmr1* altered synaptic event kinetics without reproducing the adaptation index phenotype observed in gKO mice, we hypothesized that enhanced calcium entry through excitatory synapses, rather than intrinsic calcium conductances, may contribute to the aKO phenotype. To test this possibility, we varied both NMDA receptor decay kinetics and the calcium permeability fraction of NMDA receptors within the model. Increasing either parameter produced firing behaviors similar to those observed experimentally and recapitulated the effects of increasing synaptic conductance alone (Figure 5G, bottom panel). Together, these findings identify prolonged excitatory synaptic signaling as a major consequence of astrocytic FMRP loss and suggest that altered AMPA- and NMDA-mediated transmission contributes to the impaired firing properties of spinal PVNs. Computational modeling further indicates that enhanced synaptic calcium influx provides a plausible mechanism linking astrocyte dysfunction to altered PVN excitability.

**Table 2.** Synaptic Current Parameters.

| Parameter | Value | Unit | Definition |
| --- | --- | --- | --- |
| $\bar{g}_{NMDA}$ | 20 | nS | Maximum NMDAR conductance |
| $\bar{g}_{AMPA}$ | 10 | nS | Maximum AMPAR conductance |
| $\tau_{NMDA,r}$ | 5 | ms | NMDA rise time constant |
| $\tau_{NMDA,d}$ | 50 | ms | NMDA decay time constant |
| $\tau_{AMPA,r}$ | 0.2 | ms | AMPA rise time constant |
| $\tau_{AMPA,d}$ | 2 | ms | AMPA decay time constant |
| $P_{NMDA}^f$ | 0.1 | unitless | Ca<sup>2+</sup> permeability of NMDA |
| $P_{AMPA}^f$ | 0.03 | unitless | Ca<sup>2+</sup> permeability of AMPA |
| $V_{syn}$ | 0 | mV | Synaptic reversal potential |
| $[Mg^{2+}]_{ext}$ | 1.0 | mM | External Magnesium concentration |

## DISCUSSION

In this study, we identified the spinal dorsal horn as an additional site of circuit pathology in FXS. Specifically, we demonstrated that dorsal horn PVNs, an inhibitory population involved in gating tactile information, were lower in number and had reduced intrinsic excitability in a global *Fmr1* KO mouse model of FXS. Furthermore, we showed that these deficits are predominantly through non-cell-intrinsic mechanisms and can be largely reproduced by selective deletion of *Fmr1* in astrocytes. Finally, computational modeling showed that simulations of PVN firing in the different phenotypes requires an additional extrinsic synaptic component. Together, these findings showed that the loss of FMRP impacts spinal PVN number and their excitability through astrocyte-dependent mechanisms.

A main observation in this study is that the number of spinal PVNs in gKO are lower compared to their WT littermate controls (Figure 1). We examined multiple postnatal timepoints and found that PVNs were consistently lower in the gKO mice, indicating they did not die in adulthood, and may have failed to mature properly. PV expression is widely used as a marker of mature fast-spiking inhibitory interneurons, but its emergence is developmentally regulated and depends on neuronal activity, synaptic input, and local circuit environment^28,42,43^. In cortical circuits, PVNs undergo prolonged postnatal maturation involving changes in gene expression, synaptic connectivity, firing properties, and extracellular matrix organization^44,45^. Although the developmental factors governing cortical PVNs have been studied extensively, much less is known about the maturation of spinal PVNs. Our data showed that PVNs do not appear in the lamina II-III until after postnatal day 12, compared to their cortical counterparts which express PV at postnatal day 10^15,43,46^. Thus, the mechanisms that generate and mature spinal inhibitory interneurons are likely distinct from those in cortex. Cortical PVNs derive primarily from the medial ganglionic eminence and migrate tangentially before integrating into cortical circuits^47^, whereas spinal inhibitory interneurons arise from local dorsal progenitor cells and integrate into lamina-specific circuits^26,48^. Thus, the decrease in spinal PVNs of gKO mice may reflect disruption of region-specific developmental regulations rather than the general loss of PVNs. Our results suggest a maturation defect rather than neurodegeneration. Future lineage-tracing and developmental transcriptomic experiments will be important to determine whether loss of *Fmr1* delays PVN maturation, prevents stable maintenance of PV expression, or alters the developmental trajectory of PV-lineage neurons in the spinal dorsal horn.

The reduced ability of PVNs to sustain high frequency firing (Figure 2) provides functional evidence that the cellular defects are accompanied by altered neuronal output. Previous studies have shown that activity-dependent mechanisms regulate PV expression. In cortical PVNs, sensory experience, presynaptic activity, and neurotrophic signaling can promote PV expression and neuronal maturation^42,43^, whereas reduced activity suppresses PV expression^49,50^. Our findings that spinal PVNs in gKO mice show both reduced PV expression and impaired firing is therefore consistent with the idea that PV expression and the ability to fire at high frequencies are tightly linked during inhibitory circuit development.

Our conditional deletion experiment indicates that loss of FMRP within PVNs themselves is not sufficient to explain the spinal PVN phenotype of the gKO mice (Figure 3). Previous studies have demonstrated that selective deletion of *Fmr1* in PVNs alters protein synthesis and social behavior^51^, while restoration of FMRP expression specifically in cortical PVNs is sufficient to rescue visual experience-dependent responses and improve sensory discrimination performance in gKO mice^52^. Furthermore, genetic manipulations of PVN activity have been shown to ameliorate circuit dysfunction and behavioral abnormalities in multiple models of neurodevelopmental disorders, including FXS^14,52^. Our findings suggest that the contribution of FMRP to PVN maturation and function is dependent on factors extrinsic to PVNs themselves, such as elements of their local environment.

Astrocytes emerged as strong candidate mediators of this non-PVN-intrinsic effect. Glial cells regulate neuronal maturation, synapse formation, extracellular ion balance, neurotransmitter clearance, and circuit refinement^33^. In FXS, glial dysfunction has increasingly been recognized as a contributor to disease pathophysiology^36,37^. Astrocytes are particularly relevant because they influence inhibitory synapse development and regulate PVN maturation in several brain regions^53,54^. *Fmr1* deletion in astrocytes reduced their ability to uptake glutamate, which may contribute to changes in neuronal excitability and dysregulate synaptic signalling^55,56^. In the auditory cortex, *Fmr1* deletion in astrocytes resulted in decreased inhibitory PV synapses, decreased PV levels, but did not diminish the number of PVNs^54^. Our findings that astrocyte-specific deletion of *Fmr1* reduces PV expression and impairs firing support these findings, suggesting an astrocytic mechanism regulates PVN hypofunction, but not the number of PVNs.

Our model predicts that prolonged excitatory synaptic signaling initially enhances excitatory drive onto PVNs, accounting for the transient increase in mean firing frequency observed in aKO mice (Figure 4M). Increased AMPA or NMDA receptor activation could promote an initial influx of calcium, which activates calcium-dependent potassium conductances, producing the spike-frequency adaptation observed experimentally^57^. Interestingly, prolonging AMPA currents more faithfully reproduced the experimental phenotype than comparable perturbations of NMDA currents (Figure 5G). This difference likely reflects the fundamentally different kinetics of these receptors. AMPA-mediated synaptic currents are normally brief^58,59^, but simulating the pathology by prolongating their decay time constant would substantially increase both the amplitude and duration of the excitatory drive. The resulting depolarization would increase the firing frequency early after the start of the stimulus but eventually disrupt sustained spike generation and produce fewer action potentials, and a shorter discharge duration. In contrast, NMDA receptor-mediated currents are intrinsically slow^59,60^, and extending their decay time constant produces a comparatively modest current perturbation relative to their normal kinetics. Consequently, increasing NMDA conductance modulates excitability in largely the same direction as AMPA. However, increasing calcium permeability, 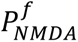 has a stronger effect than the NMDA conductance itself. Together with the conditional knockout experiments, these findings suggest that disrupted synaptic regulation, rather than intrinsic membrane dysfunction alone, is the predominant mechanism underlying impaired PVN firing following astrocytic loss of FMRP.

One difference between the model predictions and the experimental observations is that the transient increase in firing frequency was present in the aKO mice but not in gKO mice. One possible explanation is that loss of FMRP in additional cell populations, such as excitatory interneurons, primary afferents, microglia, or oligodendrocytes, engages compensatory circuit mechanisms that supress the initial increase in mean frequency while preserving the later frequency adaptation. Defining the contribution of these additional cell types will require future studies combining cell type-specific genetic manipulations with direct measurements of PVN intrinsic excitability and synaptic transmission.

In summary, our findings identify spinal dorsal horn PVNs as a previously unrecognized cellular target of loss of FMRP and demonstrate that their numbers and excitability are regulated predominantly through astrocyte-dependent mechanisms. More broadly, these results extend current models of sensory dysfunction in FXS beyond cortical and peripheral mechanisms and suggest that disrupted glia-neuron interactions within spinal sensory circuits may contribute to tactile hypersensitivity in neurodevelopmental disorders.

## MATERIALS AND METHODS

### Animals

Male mice aged 7-8 weeks old, unless otherwise stated, were kept on a 12-h:12-h light/dark cycle, with food and water provided ad libitum. All experimental procedures were approved by the Animal Care and Use Committee at McGill University, in accordance with the regulations of the Canadian Council on Animal Care. All breeders were obtained from the Jackson Laboratory and crossed in house unless otherwise stated:

1. global *Fmr1* KO mice (JAX #004624), Fmr1 WT mice (JAX # 000664);
2. PVcre;tdTomato;Fmr1^KO/y^ (gKO) and their littermate controls (JAX #008069, #007914);
3. PVcre;tdTomato;Fmr1^flox/y^ (cKO) and their littermate controls (MGI:3603442);
4. Aldh1l1^CreERT2^;Pvalb^tdTomato^; Fmr1^flox/y^ (aKO) mice and their littermate controls: Aldh1l1^CreERT2^;Pvalb^tdTomato^;Fmr1^WT/y^, Aldh1l1^WT^;Pvalb^tdTomato^;Fmr1^flox/y^, and Aldh1l1^WT^;Pvalb^tdTomato^; Fmr1^WT/y^ (JAX #031008, #027395).

#### Induction of CreERT2

Tamoxifen (Sigma #T5668; 100mg + 0.5ml EtOH 100% + 9.5ml sunflower oil (Sigma), sonicated 45 min, and stored at −20°C for a maximum of 1 month) was prepared in a 10 mg/ml solution and injected IP once daily for 3 days at 75mg/kg starting at postnatal day 10. All male mice of the litter received the same tamoxifen treatment regardless of genotype.

### Fluorescent *in situ* hybridization

Mouse lumbar spinal cords were extracted following an intraperitoneal injection of 2,2,2-Tribromoethanol (Avertin, 250mg/kg) for deep anesthesia. Then, the tissue was embedded in Optimal Cutting Temperature compound (OCT) and flash frozen on dry ice. Transverse sections (14 µm) were cut using the cryostat (Lecia Microsystems), directly mounted on Fisherbrand Superfrost Plus Microscope Slides (Cat. No. 12-550-15) and stored on dry ice before starting the RNAscope in situ hybridization. Advanced Cell Diagnostics RNAscope® Multiplex Fluorescent Reagent Kit V2 (Cat. No. 323100) was used in combination with Opal fluorophores from Akoya Biosciences with *Pvalb* (Probe-Mm Pvalb-C3, Cat. No. 421931-C3) and probe diluent (Cat. No. 300041). Control slides were run simultaneously. The signal was revealed by HRP using Opal™ 570 Reagent Pack (Cat. No. FP1488001KT). Finally, the sections were stained with DAPI and mounted using ProLong Gold antifade reagent (Invitrogen, Cat. No. P36930).

### Quantitative PCR

For the single cell-qPCR, visually identified cells were extracted from acute spinal cord slices (see Electrophysiology section) and stored in lysis buffer (Takara, 635013) in -80° C before use. mRNA extracted from single cells were converted to cDNA using SuperScript™ IV VILO™ Master Mix (Thermo Fisher Scientific – Invitrogen, 11756050). For each sample, qPCR reactions were performed in triplicates on Applied Biosystems Step One Plus Real Time qPCR System using TaqMan FastAdvanced Master Mix (Thermo Fisher Scientific - Applied Biosystems, 4444557) and Taqman Gene expression Assays (Thermo Fisher Scientific – Applied Biosystems) for the following genes: *Pvalb* (Assay ID: Mm00443100_m1), *Fmr1* (Assay ID: Mm01339582_m1). Gene expression levels were quantified by the double delta Ct analysis using *Actb* (Assay ID: Mm00607939_s1) as housekeeping reference genes.

### Immunohistochemistry

Mice were anesthetized with 5% isofluorane and perfused transcardially with 0.1 M saline phosphate buffer (PBS; [in mM] 154 NaCl, 13 Na2HPO4, 2.5 NaH2PO4, pH 7.4) followed by with 4% paraformaldehyde (PFA) in PBS. Spinal cords were then extracted by laminectomy, postfixed for 2 hours in same fixative, and cryoprotected in 30% sucrose / PBS solution at 4°C. Lumbar spinal cord sections were cut using a cryostat (Leica Microsystems). Transverse sections (25 μm-thick) were cut and placed in a 24-well plate containing PBS and stored at 4°C. Tissue sections were washed three times in PBS/0.3% Triton X-100 (PBS-T; Sigma, St. Louis, MO), and incubated in 10% normal donkey serum/PBS/0.3% Triton X-100 (NDST) for 1 hour, before adding the primary antibodies over 48 hours at 4°C in a 1% NDST solution. Sections were then washed three times with PBS-T solution and incubated for 1 hour with Alexa fluorophore-conjugated secondary antibodies at room temperature (RT). The sections were washed three times with PBS, air-dried, and coverslipped using Aqua Polymont (Polysciences, Inc., Warrington, USA). The slides were stored protected from light at 4°C until analysis.

Primary antibodies were used at the indicated concentrations: anti-parvalbumin rabbit (1:1000; Swant #PV27), anti-Pax2 goat (1:1000, R&D Systems #AF3364), anti-GFAP rabbit (1:1000, Agilent Technologies #Z033429-2), anti-GFAP rat (1:1000, Fisher #13-0300), anti-GLAST guinea pig (1:1000, Cedarlane #250114 S4), anti-GLT-1 rabbit (1:1000, Alomone #AGC-022). All primary staining were revealed with corresponding secondary antibodies conjugated to Alexa fluorophores, diluted at 1:500 in 1% NDST. All sections were incubated with NucBlue Fixed Cell Stain (DAPI) for 5 minutes at RT before mounting on microscope slides.

### Microscopy and Image Analyses

Images for quantitative analysis were acquired using a Zeiss LSM780 scanning confocal microscope. All imaging was performed on 20x/0.40LD Plan-Neofluar objective lens at 1024X1024 pixels/frame, 0.6 digital zoom and 12-bit using a 32 GaAsp detecter array. The lasers were: laser diode 405nm 30mW, Ar Ion laser 458/488/514nm 25mW, DPSS-laser 561nm 20mW, HeNe Green 543nm 1mW, HeNe RED 633nm 5mW. Image files were processed for analysis in ImageJ (NIH) using the “Blind Analysis Tools” plug-in by two independent observers. For lamina I-III region of interest (ROI) identification, anatomical dorsal horn landmarks were used. For fluorescence RNAscope quantification, an intensity threshold for each probe was set at four times the mode to define individual cells as positive or negative for mRNA expression.

### Electrophysiology

Mice were anesthetized with an intraperitoneal injection of 2,2,2-Tribromoethanol (Avertin, 250mg/kg) for deep anesthesia. The back of the mouse was shaved with an electric clipper, and a dorsal laminectomy was performed. A block containing the spinal cord was quickly transferred to a dissection dish perfused with an ice-cold (4°C)) N-Methyl-D-Glucamine-based artificial cerebrospinal fluid (NMDG-ACSF) solution (bubbled with 95% O2 and 5% CO2) containing the following (in mM): 93 NMDG, 2.5 KCl, 1.25 NaH2PO4, 30 NaHCO3, 20 HEPES, 25 glucose, 2 thiourea, 5 Na-L-ascorbate, 3 Na-pyruvate, 12 N-acetyl-L-cysteine, 0.5 CaCl2/2H2O, and 10 MgSO4/7H2O (pH 7.3–7.4 adjusted with HCl 12M) bubbled with 95% O2 and 5% CO2. The dura mater and pia were removed, and the spinal cord was secured onto an agar block using insect pins. Finally, a transverse slice (530-570 µm) of the spinal cord was obtained using a vibratome (VT1200 S; Leica). Slices were transferred to a submerged chamber containing HEPES-based recovery ACSF for 20 minutes at 34 C, equilibrated with 95% O2 and 5% CO2. Following the recovery incubation, slices were transferred to a recording chamber and continuously superfused with oxygenated ACSF (in mM): 119 NaCl, 24 NaHCO3, 2.5 KCl, 1.25 NaH2PO4, 2 CaCl2, 2 MgCl2, and 12.5 glucose (bubbled with 95% O2 and 5% CO2; pH 7.3; 300±5 mOsm measured), where they were then maintained at room temperature prior to transfer to the recording chamber.

### Targeted whole-cell patch clamp

Slices were transferred to a recording chamber and continuously superfused with oxygenated ACSF (2 mL/min). Patch pipettes were pulled from borosilicate glass capillaries (Harvard Apparatus) with a P-97 puller (Sutter Instruments). They were filled with a solution containing (in mM) 135 K-Gluconate, 6 NaCl, 2 MgCl2, 10 HEPES, 0.1 EGTA, 2 MgATP, 0.8 NaGTP (pH 7.3-7.4, adjusted with KOH; osmolarity, 300 mOsm, adjusted with sucrose) and had final tip resistances of 6–8 MΩ for whole-cell recording. Neurons were viewed by an upright microscope (Olympus) with a 40X water-immersion objective, infrared differential interference contrast (IR-DIC) and fluorescence. Recordings were made in whole cell current clamp (holding potential at - 70 mV) from identified PVNs expressing tdTomato. Data were acquired with pClamp 10.0 software (Molecular Devices) using MultiClamp 700B patch-clamp amplifier and Digidata 1440A (Molecular Devices). Recordings were low-pass filtered on-line at 1 kHz, digitized at 20 kHz and stored on a PC using pClamp software (Molecular Devices). After obtaining the whole-cell recording configuration, access resistance and membrane capacitance were calculated based on the response of a 10mV hyperpolarizing voltage step from a holding potential of –70mV. Pipette offset was zeroed, and the cell was excluded if a drift of more than 5 mV was noted.

Standardized current-clamp protocols were applied per cell recording. A gap-free 90s-long recording with no injection current (I = 0) was used to measure the resting membrane potential (mV). A 1-s-long step current injection from –200 to +200 pA (50 pA step increments) was used to measure the spike count and spiking duration. A 1-s-long ramp current injection of varied slopes from 50 to 400 pA/s was used to measure the spike threshold. A 5-minute voltage-clamp protocol at -70mV was used to record the sEPSCs. All action potential properties and intrinsic excitability analysis were performed on Python software with the toolbox, eFEL^61^.

### Computational model

#### PVN Computational Model

We adapted the previously published one compartmental Hodgkin-Huxley (HH) model of PVNs^20^. We reparametrized the model to fit our own dataset to especially capture the shape of the action potential cycle in the phase plane and population spike counts.

The HH model is given by:

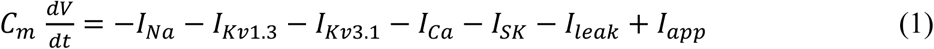

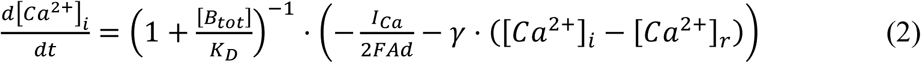

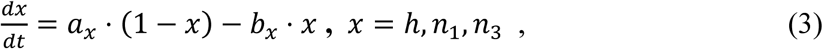

where *V* is the membrane potential of PVNs, [*Ca*^2+^]*_i_* is the intracellular calcium concentration and *x* are the ionic (activating/inactivating) gating variables.

The ionic current *I_j_* (*j=Na, Kv1.3, Kv3.1, Ca, SK*) are the various ionic currents incorporated into the model. That includes:

1. *Fast transient sodium current (I_Na_),* given by

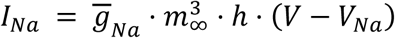

where 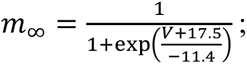
2. *Slow delayed-rectifier potassium current (I_Kv_*_1.3_), given by

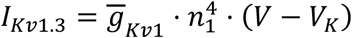
3. Fast delayed-rectifier potassium current (*I_Kv_*_3.1_), given by

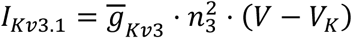
4. *High-voltage activated Calcium current (I_Ca_)*, given by;
where 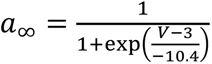
5. *Calcium-activated Potassium current (I_SK_)*, given by

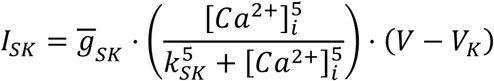
6. *Leak current (I_leak_)*, given by

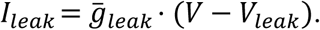

The applied current (*I_app_*) for simulations is similar in the timing and magnitude from the step currents in the voltage clamp experiments. Model parameters are provided in Table 1.

#### Modelling Synaptic Currents

To model astrocytic effects on the PVN, we added synaptic currents in addition to the HH model in Equation 1. Synaptic current were formulated from our previous work^62^. Specifically, both the AMPA and the NMDA currents were modeled using differences of exponentials as follows:

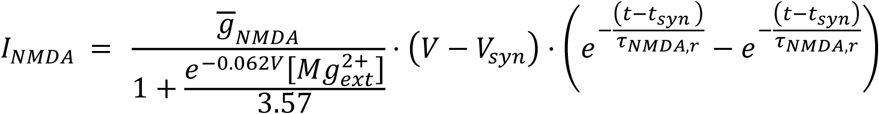

and

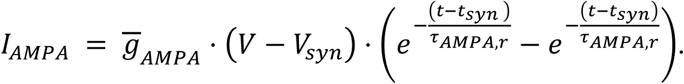

Consequently, the internal calcium concentration ([*Ca*^2+^]*_i_*) dynamics was modified as follows

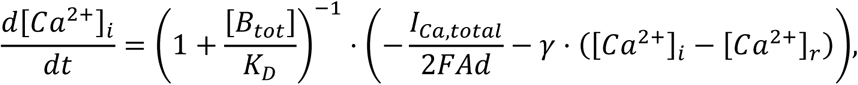

where 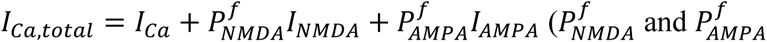 denote the fractions of the NMDA- and AMPA-mediated currents carried by calcium) and *I_syn_* = *I_NMDA_* + *I_AMPA_* is added to the right-hand side of Equation 1.

#### Feature Extraction and Parameter Estimation of the PVN model

To identify parameter sets in the seven-conductance Hodgkin-Huxley model consistent with the recorded PVN cohort, we implemented a population-level Bayesian inference pipeline. We performed different fits for each phenotype separately. A broad log-uniform prior over the maximal conductance 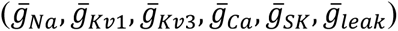 and a linear-uniform prior over the calcium buffer concentration (*B_tot_*) were sampled to generate N = 20000 candidate parameter sets. Each parameter set was simulated using the canonical 200 pA, 1 s current-injection protocol applied experimentally, and a fixed set of firing features was extracted from each simulated voltage trace using the same pipeline applied to the recordings, including spike count, adaptation index, discharge time, sag shape, and single-spike shape metrics (e.g. action potential peak, trough, and half-width, averaged over the first five spikes).

All features in Supplementary Tables 1, 2, and 3 were calculated using the Electrophys Feature Extraction Library^61,63^. For features that differed significantly between phenotypes include: Spike count (eFEL: ‘Spikecount’) is the total number of action potentials during the 1 s step current; discharge time computed as the time difference between the first and last action potential peaks (eFEL: ‘peak_time’); ean firing frequency (eFEL: mean_frequency) is defined as the number of action potentials during the stimulus divided by the time from stimulus onset to the last spike. Adaptation index, *a* (eFEL: ‘adaptation_index’) is the mean of the normalized differences between consecutive ISIs, i.e.,

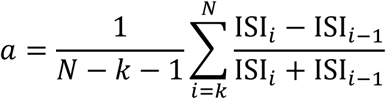

where N is the number of spikes and the first k spikes are skipped according to the eFEL default parameters. Positive values indicate spike-frequency slowing. The frequency decay time constant, *τ_f_* was computed following the approach described in ElecFeX^64^. Instantaneous firing frequencies were computed as *f_i_* = 1/ISI*_i_*, and a single-exponential decay of the form:

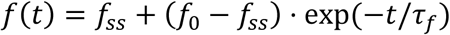

was fit to the frequency-time series.

Parameter inference was performed using Sequential Neural Posterior Estimation (SNPE)^65^, a simulation-based inference method that trains a neural density estimator to approximate the posterior *p*(*θ*|*x*) (parameter, summary statistic) samples. A masked autoregressive flow was trained on all viable simulations (i.e., simulations producing at least two spikes with all conditioning features defined), using log-transformed conductances and linear-scale *B_tot_* to match the sampling geometry. After training, the flow was conditioned on the cohort mean feature vector, and 10,000 samples were drawn from the resulting posterior distribution to characterize parameter marginals, joint correlations, and 5–95% credible intervals. To identify discrete representative parameter sets for downstream perturbation analysis, 300 posterior samples were re-simulated using the full model. The resulting simulations were then ranked according to NaN-aware root-mean-squared z-distance from the cohort mean (weighted by cohort standard error); the top 30 were retained and made sure the corresponding feature distance lay within one standard deviation from the mean of the experimental features. The same procedure was repeated on the other cohorts mean to obtain phenotype-specific posterior distributions. The best set of parameters were presented in Table 1 and Table 2.

### Software and numerical simulations

Electrophysiological features were extracted using eFEL^61,63^, and custom-written functions were used for the discharge time and frequency decay time constant. SNPE inference was performed with the sbi toolbox^65^ built on PyTorch. All analysis, statistical tests, and figures were generated in Python.

### Statistics

All statistical analyses were performed using GraphPad Prism v9.3.0 software or Python and individual test names are indicated in figure legends. Data was presented as mean ± standard error of the mean (SEM). A p-value of p < 0.05 was the criteria for significance. n.s. was denoted as not statistically significant.

## Author contributions

Conceptualization: HQ, A. Khadra, RSN Methodology: HQ, RFJ, AD, NR, CC, AV Visualization: HQ, RFJ, AD, NR, CC, AV Funding acquisition: A Khadra, RSN Writing – original draft: HQ, RFJ Writing – review & editing: all

## Classification

Biological Sciences; Neuroscience

## The Supplementary Information file includes

Supplementary Figures 1 to 3

Supplementary Tables 1 to 3

## RESOURCE AVAILABILITY

### Code accessibility

All Python scripts used to implement the computational model, perform numerical simulations, and analyze the results are publicly available on GitHub: https://github.com/rfritzdj/fmrp-ko-neuron-model. The repository includes the source code required to reproduce the computational analyses presented in this study.

### Lead Contact

Requests for further information and resources should be directed to and will be fulfilled by the lead contact, Reza Sharif-Naeini, and Anmar Khadra.

## ACKNOWLEDGEMENTS

We are thankful to the staff at the McGill University Comparative Medicine and Animal Resources Centre for the mice colony management.

