## Supplementary Information for "Astrocytic FMRP regulates the function of spinal parvalbumin-expressing neurons in Fragile X Syndrome"

**Affiliations**
1. Department of Physiology, McGill University, Montreal, QC, Canada
2. Alan Edwards Center for Research on Pain, McGill University, Montreal, QC, Canada
3. Quantitative Life Sciences, McGill University, Montreal, QC, Canada
4. Center of Applied Mathematics in Bioscience and Medicine, McGill University, Montréal, QC, Canada

† These authors contributed equally: Haoyi Qiu and Rian Fritz Jalandoni
*Corresponding authors: Haoyi Qiu, Anmar Khadra, PhD, Reza Sharif-Naeini, PhD
Senior authors: Anmar Khadra, PhD and Reza Sharif-Naeini, PhD

Mailing address:
3649 Promenade Sir William Osler
Bellini Life Sciences Center, McGill University
Montreal, Quebec, Canada H3G 1Y7

**The Supplementary Information file includes:**

Supplementary Figures 1 to 3
Supplementary Tables 1 to 3

Figure S1

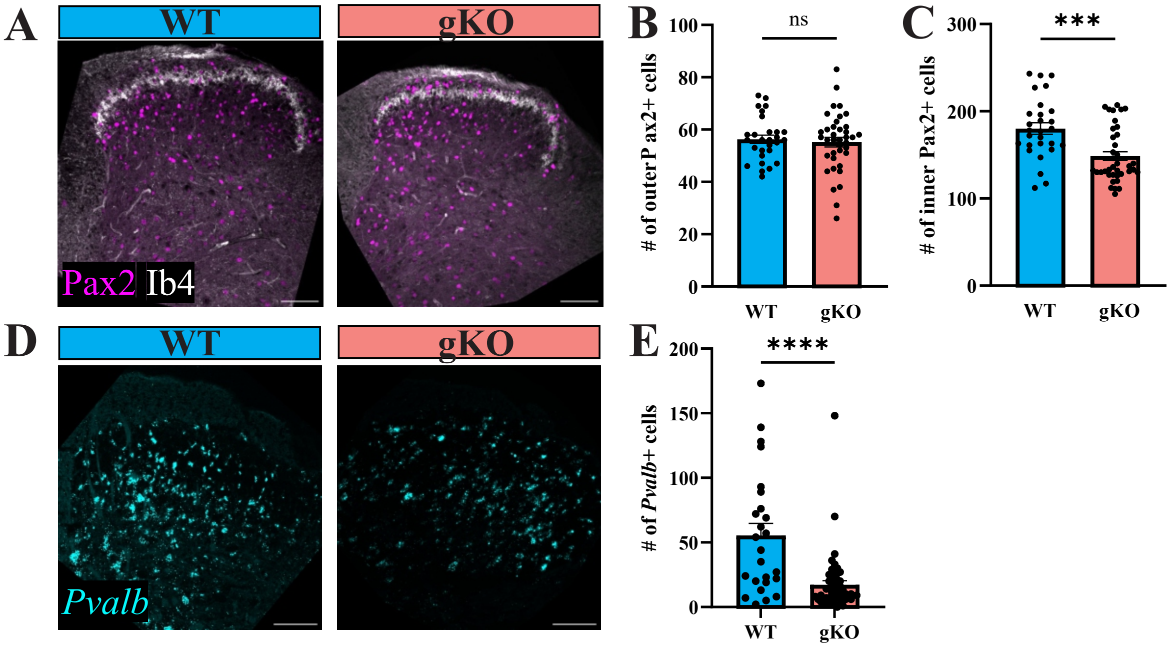

A
WT
gKO
Pax2 Ib4
B
ns
### of outer P ax2+ cells
100
80
60
40
20
0
WT
gKO
C
***
### of inner Pax2+ cells
300
200
100
0
WT
gKO
D
WT
gKO
Pvalb
E
****
### of Pvalb+ cells
200
150
100
50
0
WT
gKO

Figure S2

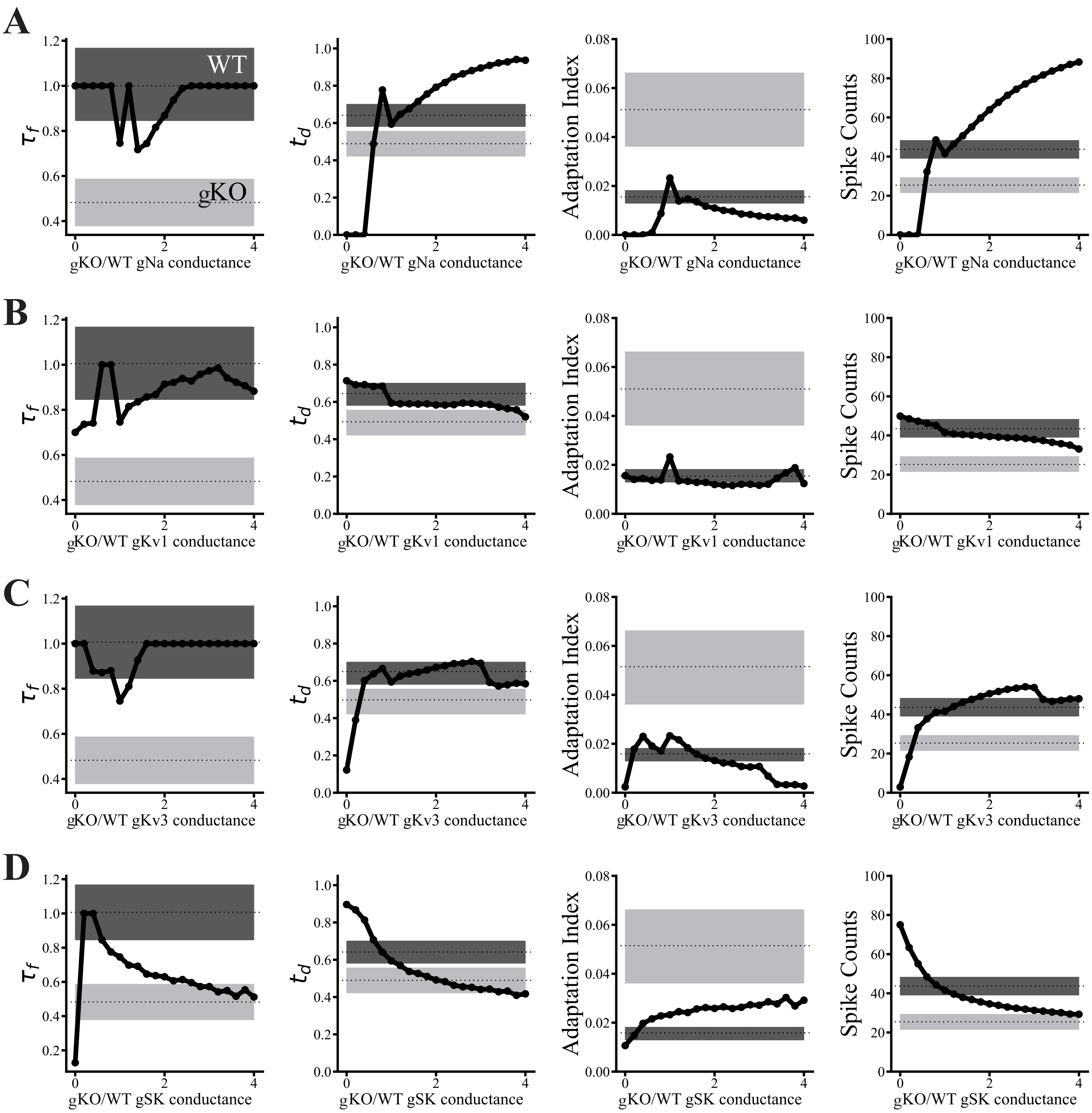
A
B
C
D
WT
gKO
$\tau_f$
$t_d$
Adaptation Index
Spike Counts
gKO/WT gNa conductance
gKO/WT gKv1 conductance
gKO/WT gKv3 conductance
gKO/WT gSK conductance

Figure S3

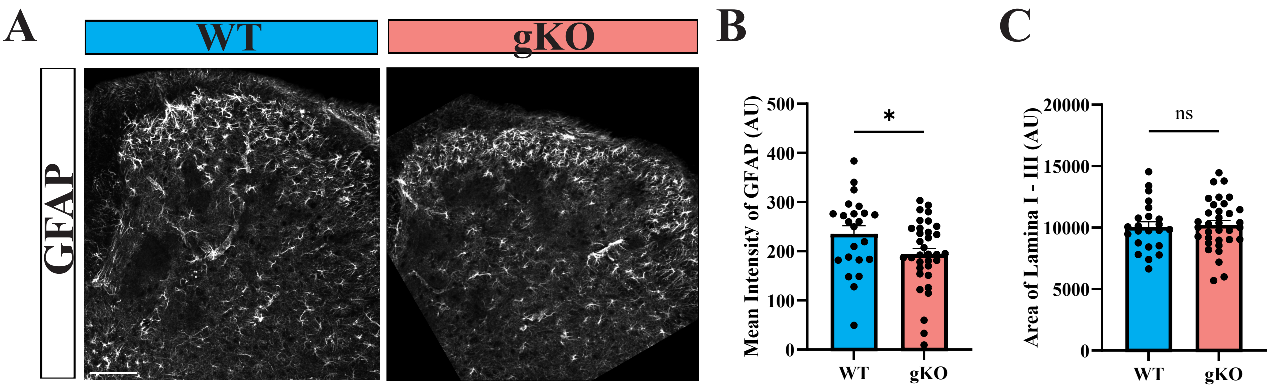
A
WT
gKO
GFAP
B
Mean Intensity of GFAP (AU)
500
400
300
200
100
0
*
WT
gKO
C
Area of Lamina I - III (AU)
20000
15000
10000
5000
0
ns
WT
gKO

Figure S1: Decreased spinal *Pvalb* expression in global *Fmr1* KO

(A) Immunohistochemistry (IHC) staining of dorsal horn inhibitory Pax2-expressing neurons (magenta), and IB4 (white) in $Fmr1^{WT/y}$ mice (WT, left) and $Fmr1^{KO/y}$ mice (gKO, right); scale bar = 100 μm.

(B-C) Mean ± S.E.M of the number of inhibitory neurons in outer (B) and inner (C) regions relative to IB4 staining in A. The outer region being from lamina I to II inner. The inner region is from lamina II inner to lamina III, which is marked by anatomical landmarks. WT: n = 28 sections from 4 male mice; gKO: n = 38 sections from 5 male mice, unpaired t-test, two-tailed. n.s. not statistically significant, ***p-value = 0.0003.

(D-E) Fluorescent *in situ* hybridization of dorsal horn (D) and quantification (E) showing the number of *Pvalb*-positive cells (cyan) in WT mice (left) and gKO mice (right); scale bar = 100μm. WT: n = 14 sections from 3 male mice; gKO: n = 26 sections from 3 male mice, unpaired t-test, two-tailed. ****p-value < 0.0001.

Figure S2: Perturbation of individual ion channel conductances fails to reproduce the global *Fmr1* KO phenotype

(A-D) Effects of varying the conductance density (gKO/WT ratio) of (A) gNa, (B) gKv1, (C) gKv3, and (D) gSK in the WT model on the frequency decay time constant (τf), discharge time (td), adaptation index, and spike count. Black curves represent simulation results across conductance values as done in Figure 2F. The experimental mean (shaded) ± S.E.M (dotted line) WT and gKO ranges are indicated in dark and light grey respectively. While varying individual conductances alters specific electrophysiological features, none reproduces the full set of changes observed in the gKO model.

Figure S3: Dysfunction in astrocytes upon *Fmr1* deletion

(A-C) Representative images (A), mean ± S.E.M of the mean intensity of GFAP (B), and the quantified area of lamina I-III (C) in WT and gKO mice. n= 23 dorsal horns from 4 WT mice, n= 34 dorsal horns from 3 gKO mice. Unpaired t-test, two-tailed, *p-value = 0.0457, scale bar = 100 μm.

**Supplementary Table 1**: Action potential properties and spiking properties of the intrinsic firing of PVNs in WT littermate and global *Fmr1* KO (gKO) mice averaged over the first five spikes or maximum available. Mann-Whitney U test.

| Action Potential Properties | WT (n=31) | gKO (n=34) | p-value |
|---|---|---|---|
| AP peak (mV) | 14.821 ± 0.892 | 13.470 ± 1.165 | 0.3477 |
| AP trough (mV) | -40.265 ± 1.400 | -40.607 ± 1.175 | 0.9948 |
| Spike threshold (mV) | -27.633 ± 1.013 | -27.988 ± 0.951 | 0.9843 |
| Spike half width (ms) | 1.874 ± 0.329 | 7.669 ± 5.242 | 0.4905 |
| AP rise time (ms) | 1.116 ± 0.036 | 1.135 ± 0.026 | 0.4341 |
| AP fall time (ms) | 1.757 ± 0.047 | 1.776 ± 0.042 | 0.9738 |
| AP rise rate (mV/ms) | 40.199 ± 1.667 | 38.164 ± 1.228 | 0.1659 |
| AP fall rate (mV/ms) | -29.395 ± 1.675 | -28.175 ± 1.221 | 0.7777 |
| AP peak to trough time (ms) | 3.146 ± 0.152 | 3.248 ± 0.148 | 0.6316 |
| AP peak to trough height (mV) | 55.086 ± 1.651 | 54.077 ± 1.246 | 0.7377 |
| Spiking Properties | | | |
| Mean Frequency (Hz) | 72.981 ± 3.782 | 77.095 ± 7.317 | 0.8284 |
| ISI CV | 0.202 ± 0.014 | 0.390 ± 0.075 | 0.4373 |
| Adaptation Index | 0.015 ± 0.003 | 0.054 ± 0.017 | **0.0028**** |
| Latency (ms) | 6.079 ± 0.348 | 6.855 ± 0.302 | 0.0630 |
| Discharge Time (s) | 0.618 ± 0.063 | 0.448 ± 0.068 | **0.0260*** |
| Sag Ratio | 0.271 ± 0.016 | 0.327 ± 0.027 | 0.1740 |
| Sag Time Constant (ms) | 101.178 ± 13.221 | 132.138 ± 21.560 | 0.5072 |
| Spike Count | 42.419 ± 4.637 | 23.647 ± 3.989 | **0.0011**** |
| Initial Frequency (Hz) | 127.143 ± 4.830 | 125.483 ± 4.512 | 1.0000 |
| Depolarized Base (mV) | -38.268 ± 1.102 | -38.736 ± 0.996 | 0.8593 |
| Frequency Decay Constant (s) | 1.065 ± 0.1613 | 0.4825 ± 0.1043 | **0.0012*** |
| Frequency Asymptote (Hz) | 54.806 ± 4.317 | 47.001 ± 4.857 | 0.1986 |

**Supplementary Table 2**: Action potential properties and spiking properties of the intrinsic firing of in PVcre;tdTomato;Fmr1$^{WT/y}$ (WT) and PVcre;tdTomato;Fmr1$^{flox/y}$ (cKO) mice averaged over the first five spikes or maximum available. Mann-Whitney U test.

| Action Potential Properties | WT (n=20) | cKO (n=21) | p-value |
|---|---|---|---|
| AP peak (mV) | 20.329 ± 1.294 | 18.936 ± 1.249 | 0.4276 |
| AP trough (mV) | -40.226 ± 1.602 | -38.305 ± 1.429 | 0.4575 |
| Spike threshold (mV) | -27.027 ± 0.868 | -26.188 ± 1.263 | 0.7914 |
| Spike half width (ms) | 1.713 ± 0.106 | 4.179 ± 1.740 | 0.8898 |
| AP rise time (ms) | 1.210 ± 0.057 | 1.195 ± 0.043 | 0.9197 |
| AP fall time (ms) | 1.897 ± 0.063 | 1.904 ± 0.060 | 0.7528 |
| AP rise rate (mV/ms) | 42.374 ± 3.025 | 40.180 ± 2.252 | 0.6777 |
| AP fall rate (mV/ms) | -30.521 ± 2.820 | -27.488 ± 1.988 | 0.5372 |
| AP peak to trough time (ms) | 3.315 ± 0.184 | 3.876 ± 0.372 | 0.6233 |
| AP peak to trough height (mV) | 60.554 ± 2.714 | 57.241 ± 2.093 | 0.3713 |
| Spiking Properties | | | |
| Mean Frequency (Hz) | 69.680 ± 5.279 | 64.401 ± 4.254 | 0.6963 |
| ISI CV | 0.277 ± 0.071 | 0.257 ± 0.046 | 0.4886 |
| Adaptation Index | 0.020 ± 0.005 | 0.014 ± 0.004 | 0.7150 |
| Latency (ms) | 6.586 ± 0.437 | 6.635 ± 0.343 | 0.7817 |
| Discharge Time (s) | 0.679 ± 0.084 | 0.633 ± 0.074 | 0.3849 |
| Sag Ratio | 0.298 ± 0.024 | 0.322 ± 0.029 | 0.4424 |
| Sag Time Constant (ms) | 143.184 ± 22.408 | 129.146 ± 23.895 | 0.2845 |
| Spike Count | 42.600 ± 5.905 | 37.773 ± 4.856 | 0.5708 |
| Initial Frequency (Hz) | 114.410 ± 4.613 | 118.184 ± 3.914 | 0.4344 |
| Depolarized Base (mV) | -37.914 ± 0.920 | -36.798 ± 1.171 | 0.5710 |
| Frequency Decay Constant (s) | 1.0219 ± 0.1613 | 0.86 ± 0.1359 | 0.7377 |
| Frequency Asymptote (Hz) | 47.300 ± 4.889 | 47.730 ± 3.414 | 0.7721 |

**Supplementary Table 3**: Action potential properties of the intrinsic firing of PVNs in WT littermate and Aldh1l1$^{CreERT2}$;Pvalb$^{tdTomato}$; Fmr1$^{flox/y}$ (aKO) mice averaged over the first five spikes or maximum available. Mann-Whitney U test.

| Action Potential Properties | WT (n=20) | aKO (n=19) | p-value |
|---|---|---|---|
| AP peak (mV) | 18.459 ± 0.957 | 14.818 ± 1.284 | 0.086 |
| AP trough (mV) | -33.635 ± 1.026 | -33.928 ± 1.551 | 0.7045 |
| Spike threshold (mV) | -22.792 ± 0.748 | -25.469 ± 0.975 | 0.0657 |
| Spike half width (ms) | 1.939 ± 0.074 | 2.128 ± 0.146 | 0.4233 |
| AP rise time (ms) | 1.301 ± 0.039 | 1.359 ± 0.056 | 0.3990 |
| AP fall time (ms) | 2.056 ± 0.054 | 61.026 ± 27.375 | 0.4820 |
| AP rise rate (mV/ms) | 32.832 ± 1.217 | 32.981 ± 2.051 | 0.5838 |
| AP fall rate (mV/ms) | -22.116 ± 0.951 | -21.078 ± 1.739 | 0.6429 |
| AP peak to trough time (ms) | 3.901 ± 0.135 | 4.079 ± 0.239 | 0.8550 |
| AP peak to trough height (mV) | 52.094 ± 1.748 | 48.746 ± 2.273 | 0.2551 |
| Spiking Properties | | | |
| Mean Frequency (Hz) | 59.313 ± 3.034 | 80.078 ± 5.156 | **0.0017**** |
| ISI CV | 0.215 ± 0.022 | 0.239 ± 0.062 | 0.3008 |
| Adaptation Index | 0.012 ± 0.002 | 0.051 ± 0.018 | 0.0772 |
| Latency (ms) | 6.750 ± 0.265 | 5.947 ± 0.431 | **0.0491*** |
| Discharge Time (s) | 0.734 ± 0.057 | 0.400 ± 0.100 | **0.0073**** |
| Sag Ratio | 0.272 ± 0.019 | 0.354 ± 0.049 | 0.3915 |
| Sag Time Constant (ms) | 139.437 ± 22.698 | 235.246 ± 52.075 | 0.5838 |
| Spike Count | 45.000 ± 4.926 | 27.474 ± 6.769 | **0.0212**** |
| Initial Frequency (Hz) | 112.123 ± 6.175 | 114.251 ± 5.724 | 0.7042 |
| Depolarized Base (mV) | -32.520 ± 0.901 | -34.898 ± 1.170 | 0.0851 |
| Frequency Time Constant (s) | 1.123 ± 0.1320 | 0.7263 ± 0.1837 | **0.0466*** |
| Frequency Asymptote (Hz) | 46.391 ± 3.625 | 48.946 ± 5.833 | 0.3477 |
